# Targeting riboswitches with synthetic small RNAs for metabolic engineering

**DOI:** 10.1101/2021.06.21.449321

**Authors:** Milca Rachel da Costa Ribeiro Lins, Laura Araujo da Silva Amorim, Graciely Gomes Correa, Bruno Willian Picão, Matthias Mack, Marcel Otávio Cerri, Danielle Biscaro Pedrolli

## Abstract

Our growing knowledge of the diversity of non-coding RNAs in natural systems and our deepening knowledge of RNA folding and function have fomented the rational design of RNA regulators. Based on that knowledge, we designed and implemented a small RNA (sRNA) tool to target bacterial riboswitches and activate gene expression. The synthetic sRNA is suitable for regulation of gene expression both in cell-free and in cellular systems. It targets riboswitches to promote the antitermination folding regardless the cognate metabolite concentration. Therefore, it prevents transcription termination increasing gene expression up to 103-fold. We successfully used sRNA arrays for multiplex targeting of riboswitches. Finally, we used the synthetic sRNA to engineer an improved riboflavin producer strain. The easiness to design and construct, and the fact that the riboswitch-targeting sRNA works as a single genome copy, make it an attractive tool for engineering industrial metabolite-producing strains.

**Graphical Abstract:** 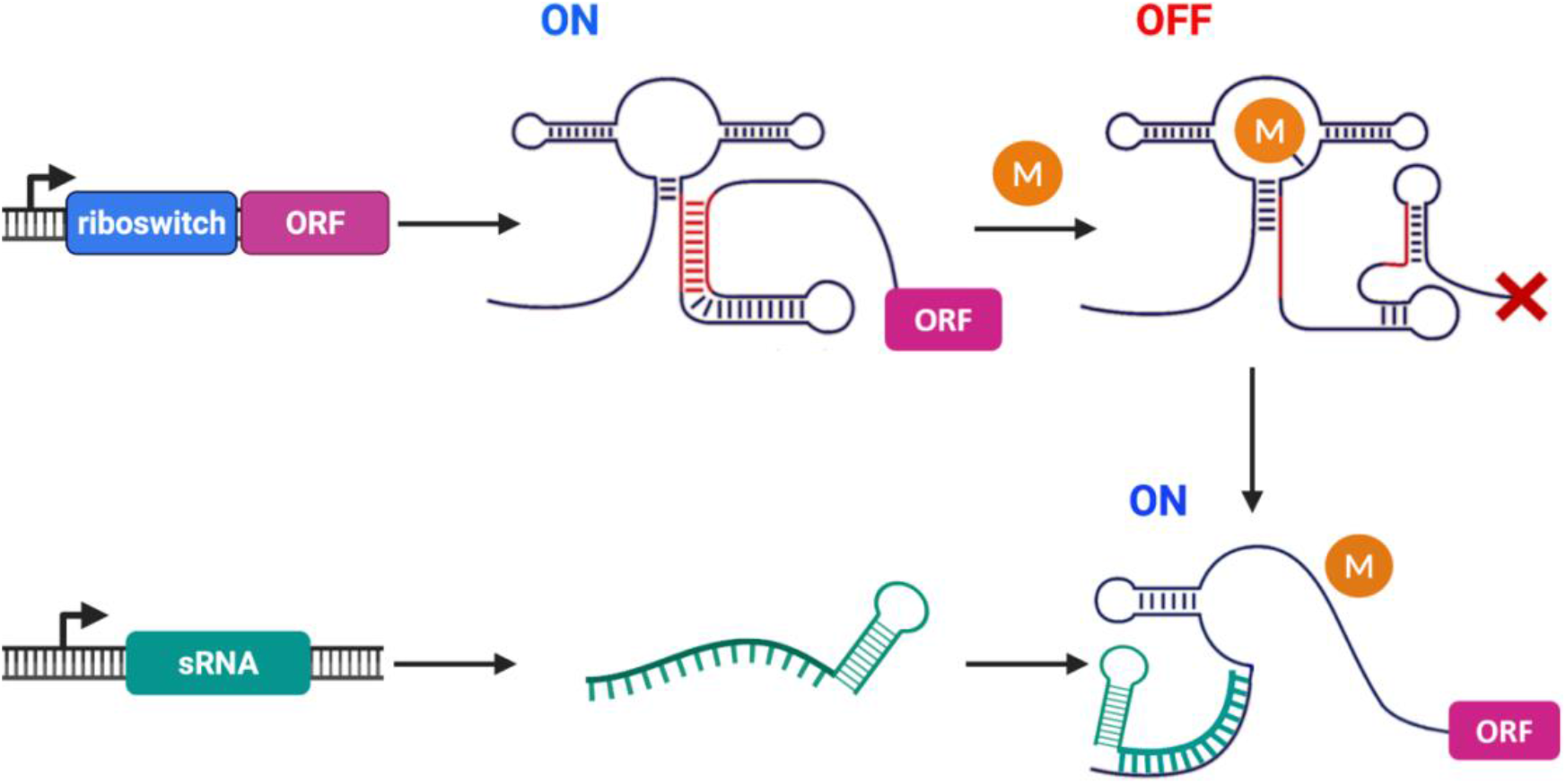

## 1. Introduction

RNAs are important regulatory tools in the synthetic biology toolbox for controlling gene expression and constructing synthetic gene networks (Chappell et al., 2015b; Leistra et al., 2019). Among all regulatory RNAs, trans-acting antisense small RNAs (sRNAs) are particularly attractive to engineer due to their simplicity, fast response to external signals, and usefulness for fine-tuning of gene expression (Shimoni et al., 2007). The simplicity of the RNA sequence and the predictability of RNA folding has powered the development of sRNA rational design to tune gene expression. Moreover, the specificity of base-pairing and easiness to construct make sRNA an ideal tool for circuit design and multiplex control (Kelly et al., 2018; Noh et al., 2017). Compared to protein-based regulation, sRNAs offer multiple options for the design of orthogonal control with minimal or no metabolic burden associated. Therefore, synthetic sRNAs are simple and versatile tools to regulate gene expression and engineer metabolic pathways.

Synthetic sRNAs have been mostly used as antisense RNAs to block the access of the ribosome to the mRNA and prevent translation in *E. coli*. sRNA-mediated knockdown has been successfully used to fine-tune gene expression and develop *E. coli* superproducers of industrially relevant compounds such as tyrosine, proline, cadaverine, putrescine, 4-hydroxycoumarin, resveratrol, and naringenin (Na et al., 2013; Noh et al., 2017; Yang et al., 2015). The strategy has been once used in *B. subtilis* to engineer a N-acetylglucosamine producer strain (Liu et al., 2014). Now, we expand the functionality of sRNAs to target riboswitches and activate gene expression at the transcription level. Riboswitches play an important role in the regulation of gene expression in bacteria. In *B. subtilis* there are 41 identified riboswitches that regulate ∼2% of all genes, many of them related to the biosynthesis of industrially relevant compounds (Kalvari et al., 2021; Mandal et al., 2003). Bacterial riboswitches ultimately regulate the levels of metabolites in the cell through the control of biosynthesis and/or transport processes (Mandal et al., 2003; Mars et al., 2016). Although tempting, deletion of riboswitches aiming to create constitutive expression leads to severe decrease in gene expression (Boumezbeur et al., 2020; Shi et al., 2014). Alternatively, we propose the use of riboswitch-targeting sRNAs for dynamic control of gene expression.

Riboswitches that bind organic small molecules, such as purines, vitamins, and amino acids, keep the cell homeostasis preventing unneeded accumulation of cellular compounds. In *B. subtilis* and other bacteria the control exerted by riboswitches is most pronounced during the exponential growth phase when the intense cell metabolism favors biosynthetic processes (Lins et al., 2021; Pedrolli et al., 2015, 2012). The developed riboswitch-targeting sRNAs act at that time point, interfering with the riboswitch aptamer folding, preventing transcription termination, and stabilizing the mRNA, which leads to metabolite accumulation.

## 2. Materials and methods

### 2.1. sRNA design analysis *in silico*

The synthetic RNAs were designed for each riboswitch using the RiboMaker software (Rodrigo and Jaramillo, 2014; Rostain et al., 2015). The thermodynamic properties of the sRNAs were evaluated using the NUPACK Web Application (Zadeh et al., 2011).

### 2.2. Plasmids and other DNA sequences

Plasmids for riboswitch control of gene expression have been constructed previously (Lins et al., 2021). Shortly, for *in vitro* gene expression each riboswitch was cloned between the *E. coli ribB* promoter and the firefly luciferase gene (*luc*) to generate plasmid series P_*ribB*_-riboswitch-*luc*. For *in vivo* gene expression, expression each riboswitch was cloned between the P_*srfA*_ promoter and the *luxABCDE* operon to generate the plasmid series pBS3C-riboswitch-*lux*. The plasmids integrate at the *sacA* locus in *B. subtilis*.

The synthetic sRNAs were purchased as oligonucleotides forward and reverse strands, annealed, and cloned into the pBS2E plasmid (Radeck et al., 2013) together with the P_*grac*_ promoter to generate the plasmid series pBS2E-P_grac_-sRNA. The plasmids integrate at the *lacA* locus in *B. subtilis*. The P_*grac*_-sRNA*-purE*-broccoli sequence was cloned into the pBS1C plasmid (Radeck et al., 2013) and integrated at the *amyE* locus in *B. subtilis*. The sRNA operon was assembled from 28 semi-complementary oligonucleotides designed with GeneDesign (Richardson et al., 2006). Four building blocks were generated by PCR gene synthesis (Rouillard et al., 2004). The four blocks were then assembled using Overlap Extension PCR. Lists of plasmids and strains used are provided in the supplementary tables S1 e S2.

### 2.3. *In vitro* transcription

RNAs transcripts were generated by *in vitro* transcription using the RiboMAX™ Large Scale RNA Production Systems (Promega). Annealed oligonucleotides forward and reverse containing the T7 promoter and the sRNA were used as template. Reactions were incubated at 37ºC overnight. Reaction inactivation was performed at 70°C for 20 min. To eliminate the template, all samples were treated with RQ1 RNase-Free DNase (Promega) for 1 hour at 37ºC. Samples were then precipitated with ethanol and resuspended in RNase-free water.

### 2.4. *In vitro* gene expression

The regulatory activity of the sRNAs was accessed by *in vitro* gene expression reaction according to Lins et al. (Lins et al., 2021). Each P_*ribB*_-riboswitch-*luc* plasmid was extracted from overnight grown *E. coli* with a miniprep kit (PureYield(tm) Plasmid Miniprep System, Promega) and purified using ethanol precipitation. Plasmids were resuspended in RNase-free water and used as templates for *in vitro* gene expression (*E. coli* S30 Extract System for Circular DNA, Promega). Reactions were set up by adding 1 μL of 0.25 mM amino acid mix, 4 μL of S30 Premix, 3 μL of S30 Extract, 1 μL of 50 µM ligand (guanine, adenine or FMN; Merck), 1 μL of transcribed sRNA (30 ng μL^-1^), and 1 μL of plasmid (5 ng μL^-1^). The reaction mixture was incubated at 30°C for 20 min, and 90 μL of 1% (m/v) bovine serum albumin was added to stop the reactions. Each treatment was performed in triplicate.

In order to quantify the luciferase activity, 10 μL of stopped reactions were added to 50 µL of luciferase substrate solution (Luciferase Assay System kit, Promega) in a 96-well white flat bottom microplate. Luminescence readings were taken on a microplate reader (Tecan Infinite 200 Pro, Switzerland) set with 1,000 ms for integration time at 26ºC.

### 2.5. Bacterial strains and media

*E. coli* Top10 was used for cloning and propagation of plasmids. It was aerobically cultivated at 37°C in Lysogeny Broth (LB) enriched with antibiotics when needed. *B. subtilis* 168, *B. subtilis* riboflavin producer (BsRF), and *B. subtilis* Ai (BsAi) (Correa et al., 2020) were used as chassis for the experiments as indicated in the main text. All *B. subtilis* strains were aerobically cultivated in LB for all experiments regarding bioluminescence measurements. A high sugar complex medium was used for riboflavin production in all scales (PW medium: 40 g L^-1^ sucrose, 1 g L^-1^ yeast extract, 25 g L^-1^ NaNO_3_, 0.333 g L^-1^, KH_2_PO_4_, 1 g L^-1^ Na_2_HPO_4_·12H_2_O, 0.15 g L^-1^ MgSO4·7H_2_O, 7.5 mg L^-1^, CaCl_2_, 6 mg L^-1^ MnSO4·H_2_O, 6 mg L^-1^, FeSO_4_ 7H_2_O, pH 7.0). *B. subtilis* culture medium was enriched with chloramphenicol (5 µg mL^-1^) and/or erythromycin (1 µg mL^-1^) when needed.

### 2.6. Bioluminescence and growth measurements

Bioluminescence measurements from growing *B. subtilis* were carried out in white 96-well microplates with optically clear bottoms. Culture density was monitored by absorbance measurements at 600 nm (OD_600_) taken from the same transparent bottom microplates. Plates were incubated at 37ºC and orbital shaking at 143 rpm. Measurements were performed every 10 min for five hours in the microplate reader (Tecan Infinite 200 Pro). All experiments were performed with three biological replicates.

### 2.7. Broccoli-DFHBI fluorescence measurements

The RNA-Broccoli (Filonov et al., 2014) was fused to the *purE*-sRNA under control of the P_*grac*_ for detection of transcripts in *B. subtilis*. After induction with IPTG, samples were withdrawn and incubated for 30 min with the fluorophore DFHBI. Then, fluorescence measurements were taken in a 96-well black microplate with the microplate reader set at 472 nm for excitation and 507 nm for emission.

### 2.8. Test tube and Erlenmeyer scale cultivation

*B. subtilis* cultivation was performed in 16 × 220 mm (d x L) test tubes filled with 5 mL culture medium and incubated at 37°C and 220 rpm. *B. subtilis* culture was also carried out in 250 mL baffled Erlenmeyer flasks filled with 20 mL culture medium and incubated at 37°C and 150 rpm. Samples were periodically withdrawn for absorbance measurements at 600 nm (OD_600_) in a microplate reader (Tecan Infinite 200 Pro, Switzerland), and for riboflavin quantification by HPLC. All experiments were performed with three biological replicates.

### 2.9. Bioreactor scale cultivation

A stirred batch was carried out at 37ºC, 0,5 vvm air supply, and pH controlled at 7.0±0.1. The bioreactor 7L tank was filled with 5L of PW medium supplemented with 80 g.L^-1^ sucrose. After inoculation, samples were withdrawn periodically for sucrose, riboflavin, and biomass quantification. Biomass quantification was accessed as dry weight per liter.

### 2.10. Riboflavin and sucrose quantification

Samples withdrawn from cultures were centrifuged at 16,000 xg for 30 min for supernatant recovery. Trichloroacetic acid 1% (v/v) was added to the supernatant to precipitate proteins. The resulting samples were centrifuged at 16,000 xg for 20 min. The supernatant was then filtered using a hydrophilic PTFE 0.20 μm membrane, and the filtrate was used for analysis by high-performance liquid chromatography (HPLC; Shimadzu LC-20AD equipped with a SPD-M20A photodiode detector).

Riboflavin was analyzed using a 50 × 3 mm Poroshell 120 EC-C18 column (Agilent Technologies) in 82% solvent A (20 mM formic acid, 20 mM ammonium formate) and 18% solvent B (methanol) as mobile phase at 0.5 mL/min, in isothermal mode at 30°C. Detection was performed at 445 nm. Commercial riboflavin (Merck) was used as standard for calibration.

Sucrose was analyzed using a HPX-87P column (Aminex) in ultrapure water as mobile phase at 0.6 mL/min, in isothermal mode at 80°C.

### 2.11. Statistical analysis

Analysis of variance (ANOVA) and Tukey test were performed using the Minitab software (version 18), to compare the averages and observe the significance of the results.

## 3. Results

### 3.1. Designing small regulatory RNAs to target riboswitches

We have chosen the well-studied purine riboswitches *purE, xpt, nupG*, and *pbuE*, and the flavin riboswitch *ribDG* of *B. subtilis* as targets for sRNA design. Together they control the purine uptake and synthesis and the riboflavin synthesis through premature transcription termination (Johansen et al., 2003; Lins et al., 2021; Mandal et al., 2003). The sRNA was designed to target each riboswitch in order to prevent folding into the OFF structure that leads to premature transcription termination (Figure 1). The sRNA was designed to counteract the riboswitch activity and keep gene expression ON even at high cognate metabolite concentration by promoting the anti-terminator formation even when it shouldn’t. Target sequences in the aptamers have been previously identified through point mutation as crucial for the OFF state folding (Mandal and Breaker, 2004; Marcano-Velázquez and Batey, 2015; Mironov et al., 2002). sRNA sequences targeting each riboswitch were generated using RiboMaker (Rodrigo and Jaramillo, 2014) and analyzed using NUPACK (Zadeh et al., 2011). The best sRNAs were selected based on the hybridization free energy (lower than −18 Kcal/mol), the seed-based free energy, ensemble defect (lower than 10%), and the complex representation at equilibrium (Rostain et al., 2015). One sRNA targeting each riboswitch was selected for construction and test (Table S3).

**Figure 1.**
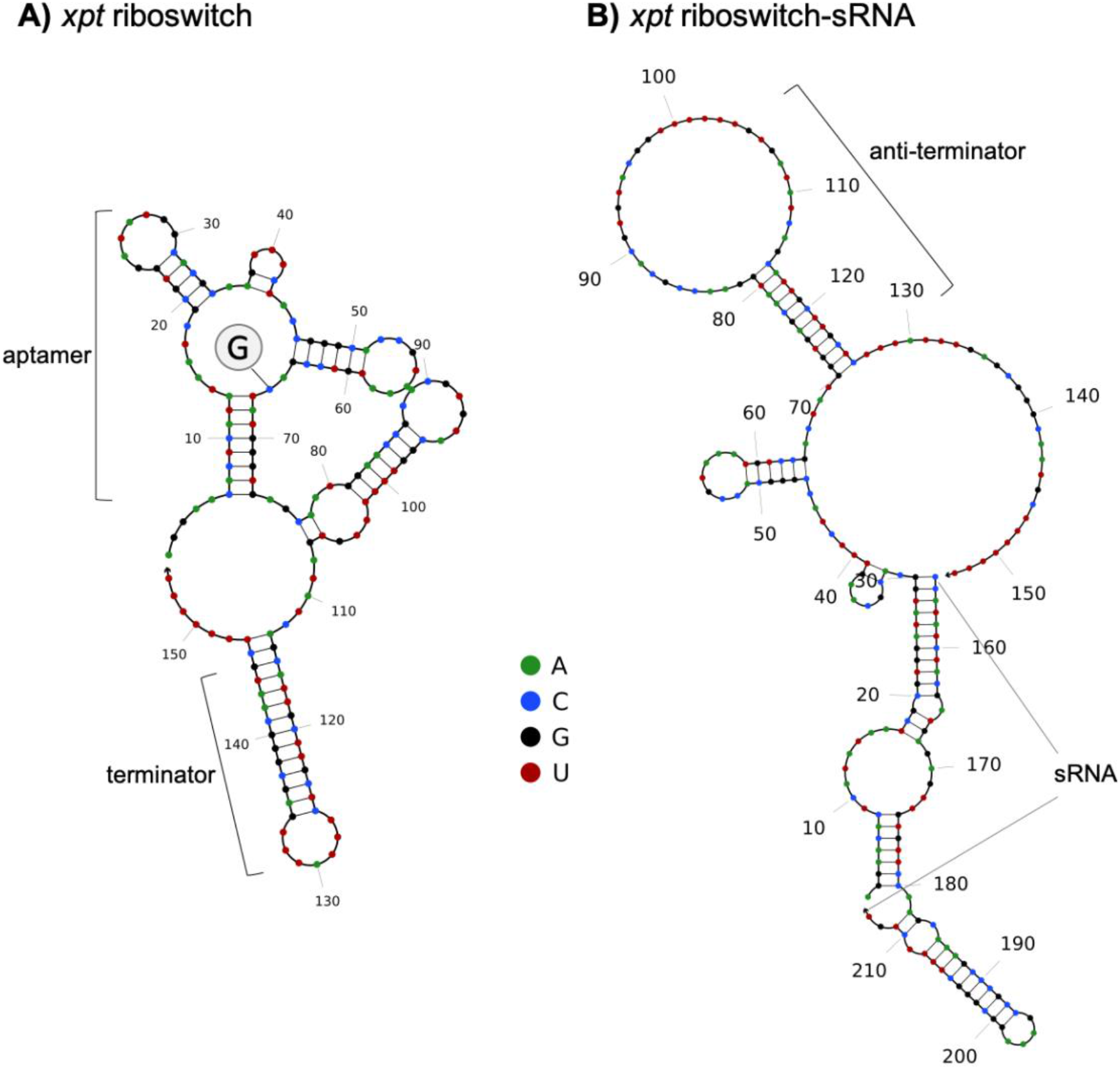
Riboswitch-sRNA paring. (A) sequence and structure of the *xpt* riboswitch bound to guanine. (B) sequence and structure of the *xpt* riboswitch paired with the *xpt*-sRNA. sRNA paring with the riboswitch nucleotides 2-30 results in an alternative folding, that leads to the formation of the anti-terminator steam loop through base-pairing between nucleotides 70-81 and 115-126. All riboswitch-targeting sRNAs were designed to target the aptamer sequence in order to promote the anti-terminator formation. Figure generated using NUPACK Web Application.

### 3.2. Synthetic sRNAs counteract riboswitches’ control of gene expression *in vitro*

In order to evaluate the specific ability of each synthetic sRNA to bind its target riboswitch and activate gene expression, we set an *in vitro* gene expression reaction which has been validated for riboswitch characterization (Lins et al., 2021). Expression of the firefly luciferase gene under the control of each riboswitch was evaluated in the presence of the sRNA and the cognate metabolite (guanine, adenine, or FMN). Each sRNA was previously transcribed *in vitro* and purified. All sRNAs were able to completely counteract their target riboswitch’s regulatory activity turning ON gene expression when it was supposed to be OFF (Figure 2). The activation fold ranged from 2.1 to 4.2. Interestingly, the sRNAs *purE, pbuE*, and *ribDG* activated gene expression to levels above the riboswitches’ ON state (Figures 2A, D, and E). The sRNA effect on the riboswitch was compared to control-sRNAs, which are riboswitch-targeting sRNAs tested against non-target riboswitches.

**Figure 2.**
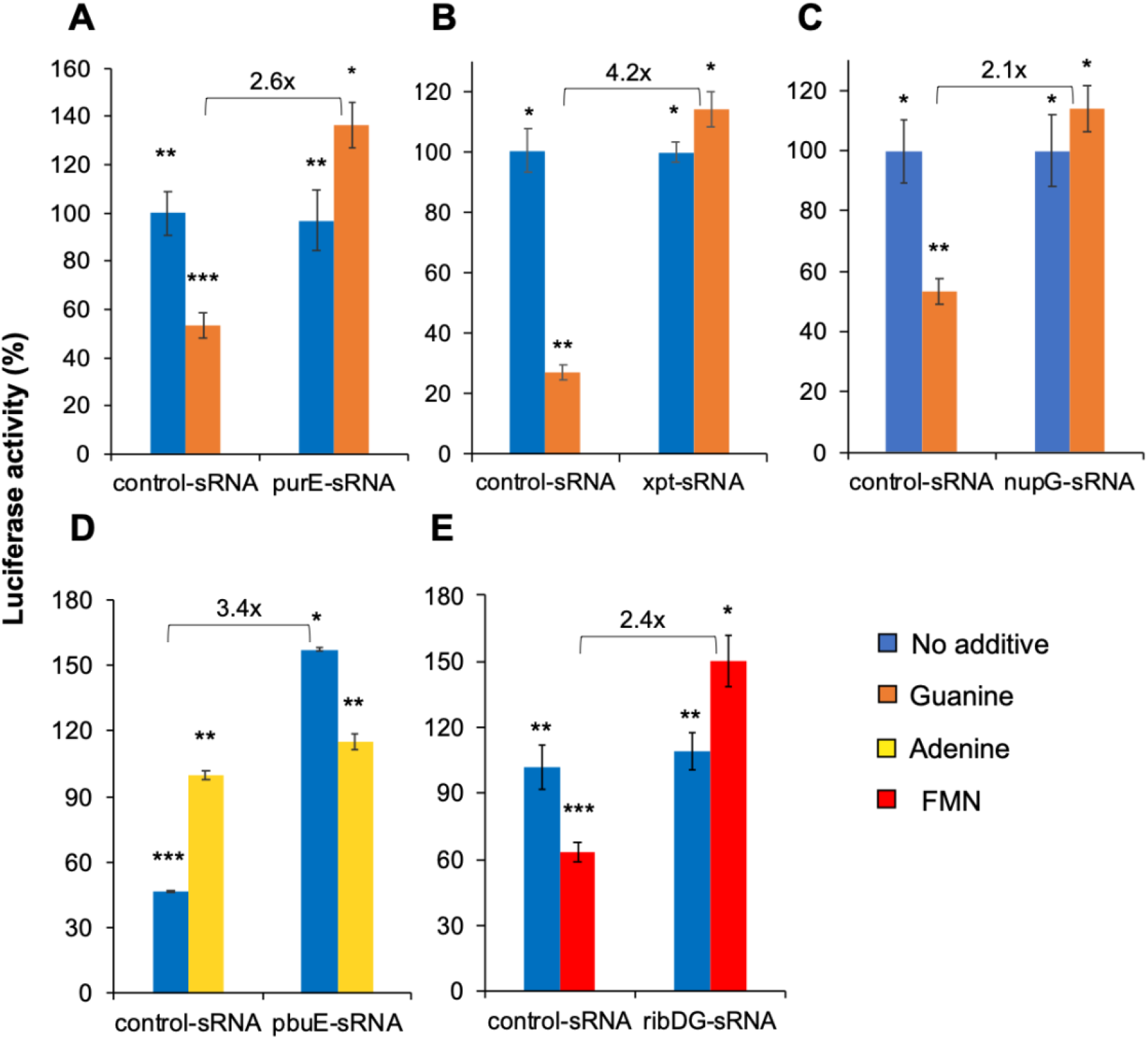
sRNA activity and specificity assayed by *in vitro* expression of the firefly luciferase gene. (A) *purE* riboswitch. (B) *xpt* riboswitch. (C) *nupG* riboswitch. (D) *pbuE* riboswitch. (E) *ribDG* riboswitch. The sRNA effect on the riboswitch was compared to control-sRNAs, which are riboswitch-targeting sRNAs tested against non-target riboswitches. Asterisks identify statistically significant differences (p < 0.05). Data are presented as mean ± SD (n = 3). Raw data available in Supplementary Data 1.

### 3.3. Synthetic sRNAs are transcribed and functional in the cell

After confirming that each synthetic sRNA can specifically counteract the target riboswitch *in vitro*, we moved to *B. subtilis* to verify whether the sRNA could be stably transcribed *in vivo*. Therefore, the *purE*-sRNA was fused to the broccoli sequence at the 3’-end and integrated as single-copy to the chromosome. Expression of the *purE*-sRNA-broccoli was induced with IPTG in the early-exponential phase, and samples were taken for DFHBI-broccoli fluorescence measurements. A fluorescence peak was detected 1h post-induction and fluorescence was still detectable after 24h (Figure 3A).

**Figure 3.**
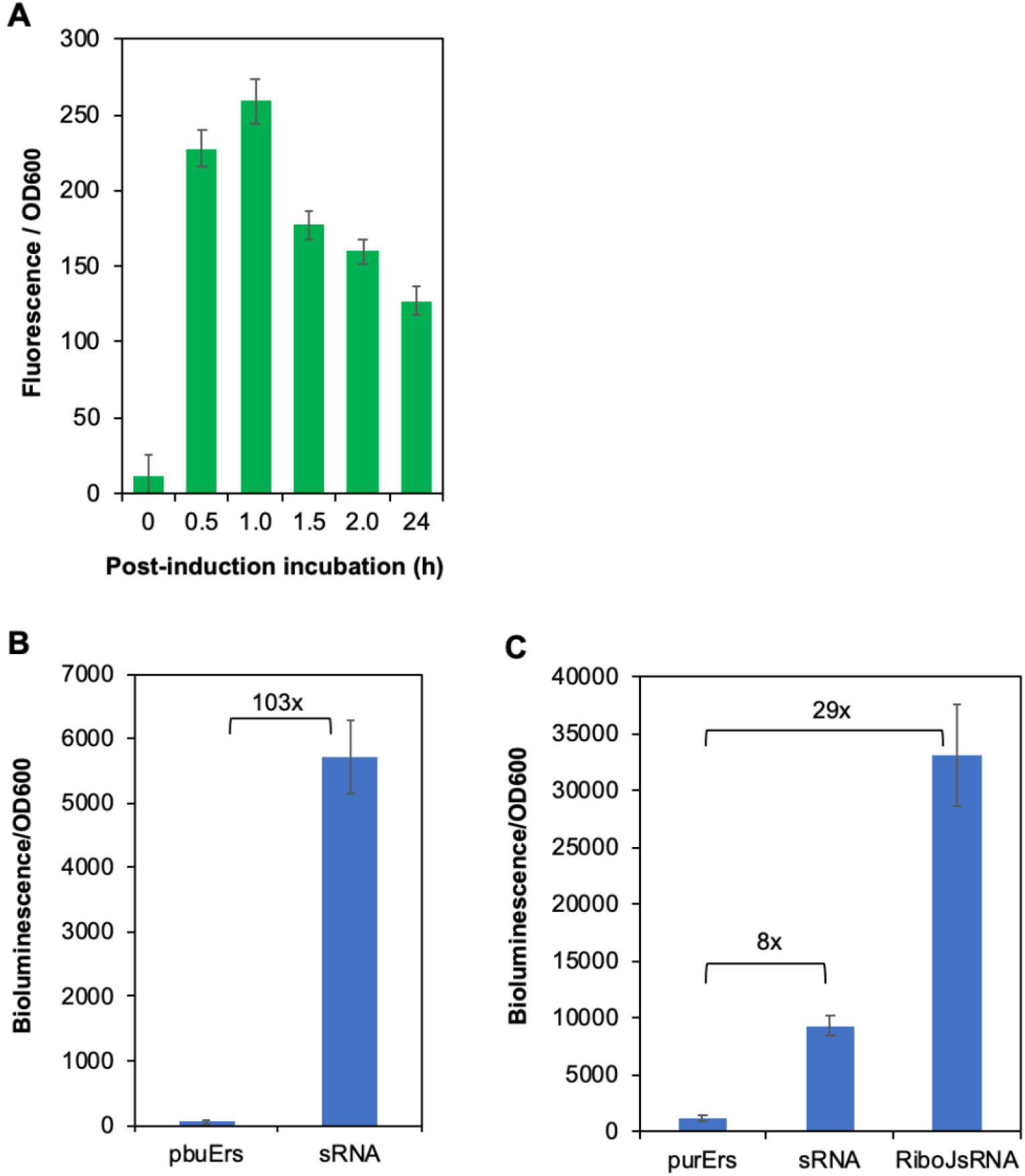
sRNA expression and activity assayed in *B. subtilis*. (A) sRNA-broccoli-DFHBI fluorescence. (B) the *lux* operon under control of the *pbuE* riboswitch (pbuErs) and the *pbuE*-sRNA (sRNA) activation effect on the riboswitch. (C) the *lux* operon under control of the *purE* riboswitch (purErs) and the *pbuE*-sRNA (sRNA) and the RiboJ-*purE*-sRNA (RiboJsRNA) activation effect on the riboswitch. Data are presented as mean ± SD (n = 3). Raw data available in Supplementary Data 2.

After validating the ability of *B. subtilis* to stably transcribe the sRNA, we constructed a reporter assay to access the sRNA activity *in vivo*. Therefore, the *luxABCDE* operon was used as a reporter under the control of the riboswitches *purE* and *pbuE*. Each riboswitch-*lux* construct was inserted as single-copy at the locus *sacA*, and the corresponding sRNA was integrated at the *lacA* locus in the chromosome. The sRNA transcription was driven by the strong P_*grac*_ promoter working constitutively (the strain lacks a *lacI* gene). The resulting strains were cultivated in Erlenmeyer flasks filled with LB medium without additives and monitored for bioluminescence for 8h. The *pbuE*-sRNA showed a strong activity activating gene expression up to 103-fold (Figure 3B), much higher than observed *in vitro*. The *purE*-sRNA caused an 8-fold increase in gene expression (Figure 3C), which is three times higher than achieved *in vitro*. Although higher than *in vitro*, gene the *purE*-sRNA activating activity *in vivo* is much lower than the activity measured for the *pbuE*-sRNA. We hypothesized that the *purE*-sRNA lower activity could be caused by low RNA stability in the cell. Therefore, we fused it to the RiboJ ribozyme to create the RiboJ*purE*-sRNA. RiboJ greatly improved the sRNA, causing a 29-fold increase in gene expression (Figure 3C). RiboJ is a structured RNA able to self-cleave at the 5’-end that has been used to insulate mRNAs (Lou et al., 2012). Any other structured RNA could have had the same beneficial effect on the sRNA; however, a self-cleaving ribozyme offers the possibility to build an sRNA array for targeting multiple loci (multiplex control). Noteworthy, the sRNAs *pbuE* and RiboJ*purE* improved the strain growth significantly compared to the control strains carrying the riboswitch-controlled *lux* operon but no sRNA (Figure S1). The low-efficient *purE*-sRNA did not cause any change in the strain growth.

### 3.4. Engineering an improved riboflavin producer strain

Synthetic sRNA has great applicability as a tool to fine-tune gene expression in metabolic pathways. However, it has mostly been used to downregulate gene expression (Liu et al., 2014; Na et al., 2013; Noh et al., 2017; Sun et al., 2019; Yang et al., 2019, 2015). In order to demonstrate the potential of the riboswitch-targeting sRNA for metabolic engineering, we tested the sRNA ability to improve the riboflavin production directly by targeting the *rib* operon, and indirectly by targeting the purine supply (Figure 4). The sRNAs *ribDG, purE, RiboJpurE, xpt*, and *pbuE* were individually inserted into the genome of a riboflavin super-producing strain (BsRF) as single copies. Riboflavin production was first evaluated in test tubes (Figure 5A). Although the sRNAs *purE* and *RiboJpurE* have been previously proved to be functional *in vivo*, they did not improve the riboflavin production. Both target the *pur* operon, which is responsible for the synthesis of inosine monophosphate (IMP) from phosphoribosyl-α-1-pyrophosphate (PRPP). IMP is the precursor for AMP, GMP, and hypoxanthine; therefore, the carbon flux may divert to other routes instead of GTP.

**Figure 4.**
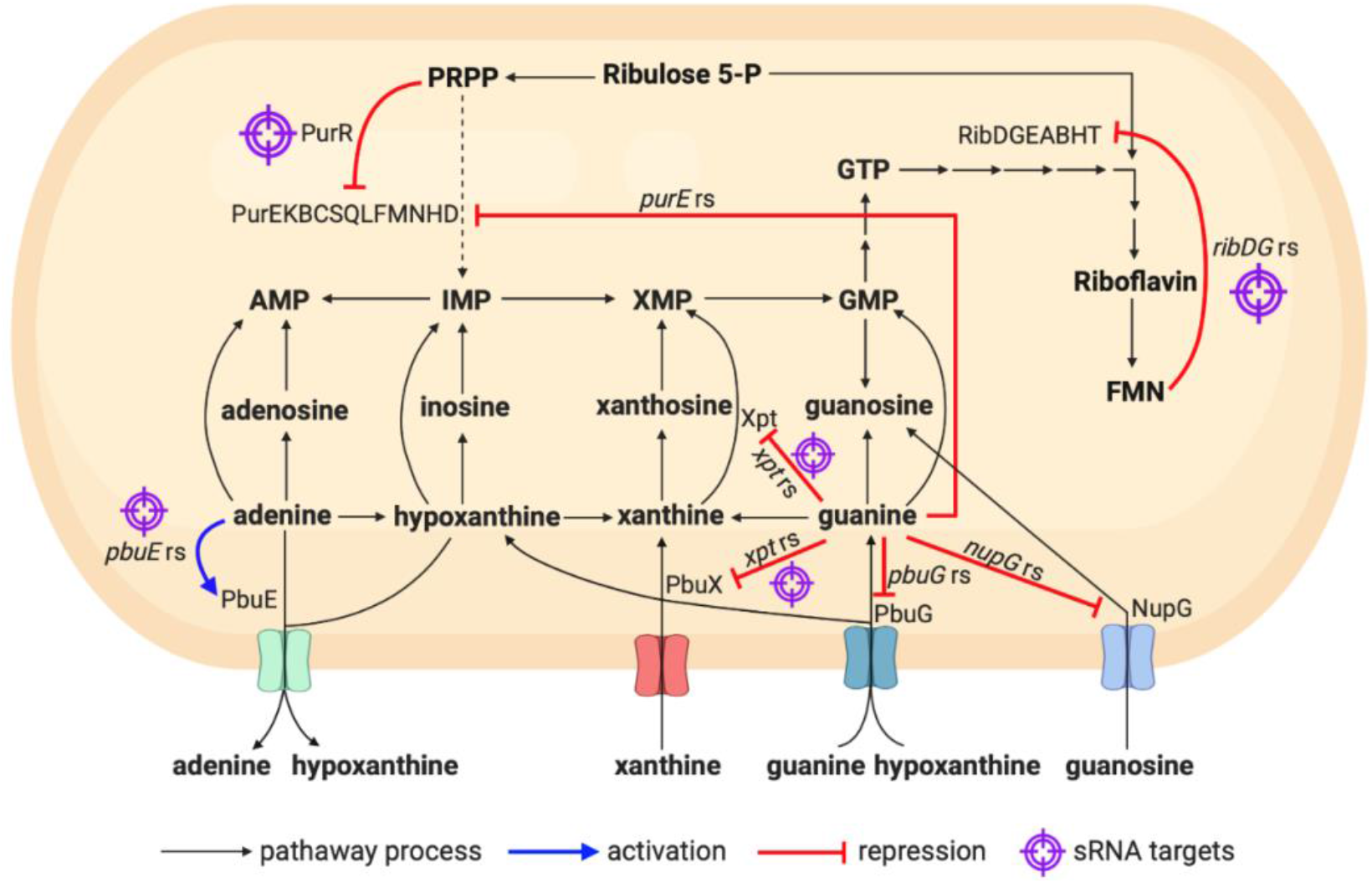
Metabolic engineering of *B. subtilis* RF (BsRF) for riboflavin production using sRNAs. The riboswitches identified as *purE* rs, *pbuE* rs, *xpt* rs, *pbuG* rs, *nupG* rs, and *ribDG* rs regulate the expression of important genes and operon from the purine and riboflavin metabolism.

**Figure 5.**
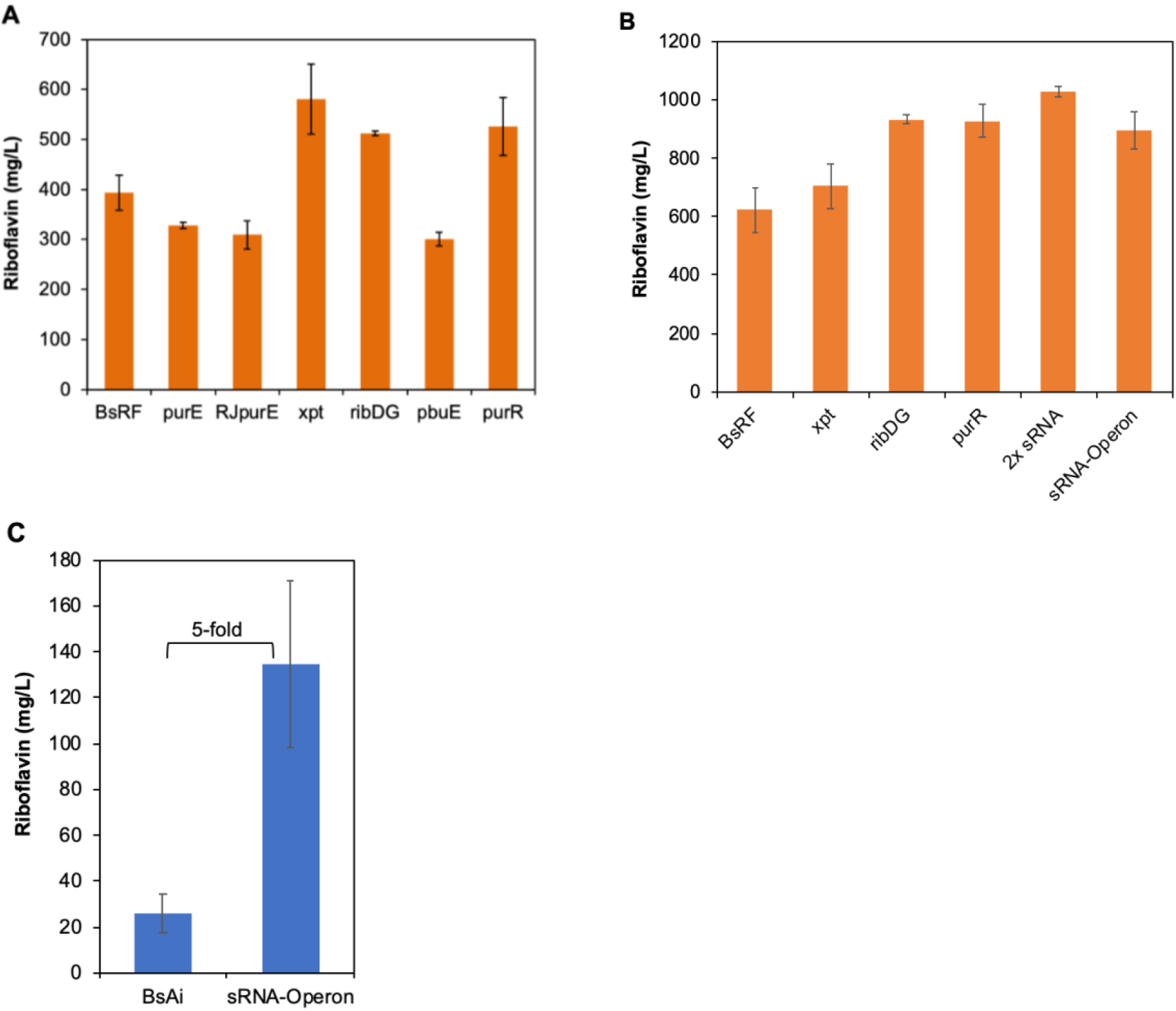
Engineering *B. subtilis* with sRNAs for riboflavin production. (A) Test tube cultivation to evaluate the effect of sRNAs on the riboflavin production. The effect of the sRNAs *purE*, RiboJ-*purE, xpt, ribDG, pbuE*, and *purR* were tested in the BsRF strain. (B) Erlenmeyer cultivation of the BsRF strains carrying the best sRNAs (*xpt, ribDG*, and *purR*) individually, an sRNA array composed of 2 sRNAs (*ribDG* and *purR*), and an sRNA operon composed of five sRNAs (*purE, xpt, ribDG, pbuE*, and *purR*). (C) The sRNA operon tested in the *B. subtilis* strain BsAi. Data are presented as mean ± SD (n = 3). Raw data available in Supplementary Data 3.

The *pbuE* sRNA targets the expression of the adenine exporter PbuE, increasing it. Although the *pbuE* sRNA has proven highly functional using the lux reporter, the strain carrying it produced lower levels of riboflavin (Figure 5A). Increased adenine efflux probably increased the IMP consumption to reset the intracellular adenine concentration, which could cause a reduced flux through GMP synthesis.

The sRNAs *xpt* and *ribDG*, on the other hand, were able to increase the riboflavin production by 48% and 30%, respectively. The *xpt* sRNA targets the expression of the *xpt*-*pbuX* operon. *pbuX* encodes a membrane protein that imports xanthine from the medium. *xpt* encodes a xanthine phosphoribosyl transferase, which converts xanthine to xanthine monophosphate (XMP) (Christiansen et al., 1997). XMP is then converted to GTP, which serves as a direct precursor to riboflavin (Figure 4). The *ribDG* sRNA targets the *ribDGBAHT* operon, which encodes the enzymes directly responsible for riboflavin biosynthesis from GTP and ribulose (Bacher et al., 1993).

To compare the performance of the designed riboswitch-targeting sRNAs with traditional antisense sRNAs (Liu et al., 2014; Na et al., 2013) we inserted the *purR*-sRNA into the *B. subtilis* chromosome. The latter was designed to pair with the *purR* mRNA from the RBS to the +14 ribonucleotide, thereby blocking translation initiation. The antisense RNA targets the synthesis of PurR, a repressor that blocks transcription initiation of several genes involved in the purine metabolism including the *pur* operon and *xpt*-*pbuX* (Saxild et al., 2001). The strain carrying the *purR*-sRNA produced 34% more riboflavin than the control (Figure 5A), resembling the *xpt*-sRNA and *ribDG*-sRNA carrying strains.

The most promising strains, carrying the sRNAs *xpt, ribDG*, and *purR*, were further tested in Erlenmeyer flasks (Figure 5B). Upscaling resulted in increased riboflavin production for all tested strains. The highest production was triggered by the sRNAs *ribDG* and *purR* reaching over 900 mg/L, which is about 50% higher than the control. The *xpt*-sRNA did not repeat the performance shown on a smaller scale and only slightly increased the riboflavin production, not significantly though. Interestingly, the strain carrying the *xpt*-sRNA grew as poor as the riboflavin super-producing strain (BsRF), whose slow growth has been ascribed to the burden of riboflavin overproduction. However, the two sRNAs that increased the riboflavin production, *ribDG* and *purR*, seem to restore the growth capability of the producer strain reaching cell densities around 2.5-fold higher than BsRF after 40h (Figure S2).

### 3.5. Multiplex sRNA for targeting riboswitches simultaneously

After testing the synthetic sRNAs individually, we wondered whether they could be combined for an additive effect on the riboflavin production. We designed a sRNA array composed of the sRNAs *purR* and *ribDG* each carrying its own promoter and terminator. Combining the two best sRNAs *purR* and *ribDG* increased 10% the riboflavin production compared to only one of them, which represents a 65% increase compared to the parental strain (Figure 5B). Next, we wanted to simultaneously test all the sRNAs developed. To avoid promoter and terminator repetition, we constructed a sRNA operon under control of a single promoter and terminator. The ribozymes RiboJ, RiboJ10, RiboJ60 e RiboJ64 (Nielsen et al., 2016) were inserted in between the sRNAs to promote the release of each sRNA after transcription. The sRNA operon was first tested in the riboflavin super-producing strain (BsRF) resulting in a 44% increase in the riboflavin titer compared to the parental strain (Figure 5B). Although functional, the sRNA operon showed a smaller effect than the best sRNAs individually, which may be attributed to a combination of the positive influence of some sRNAs (*xpt, purR*, and *ribDG*) and the negative influence of others (*purE* and *pbuE*) in the riboflavin biosynthesis. Noteworthy, both strains, the BsRF::*purR*-sRNA-*ribDG*-sRNA and the BsRF::sRNA-operon, grew to higher culture densities than the parental strain BsRF, closely reproducing the effect observed for the individual sRNAs *ribDG* and *purR*.

The BsRF strain has several mutations that increase flux through the riboflavin biosynthetic pathway and increase the availability of the precursor ribulose 6-phosphate. Therefore, we hypothesized that the sRNAs effect could be restricted in such a context. In order to evaluate the full potential of the sRNA operon, we inserted it into the genome of a *B. subtilis* strain (BsAi) that carries an extra copy of the riboflavin biosynthetic operon under control of an autoinduction device (Correa et al., 2020), but no other modification. The sRNA operon increased 5-fold the riboflavin production compared to the parental strain (Figure 5C). This result demonstrates that our synthetic sRNA operon is fully functional and capable of targeting gene expression.

### 3.6. The *BsRF*::*ribDG*-sRNA strain is stable and highly productive in a bioreactor scale

In order to demonstrate that our approach is suitable for engineering industrial metabolite-producing strains that are stable and highly productive in large scale, we scaled up the cultivation to a bioreactor. Therefore, one of our best riboflavin producing strain, the BsRF::*ribDG*-sRNA, was cultivated in a 5L batch to access its potential for large scale riboflavin production. The bioreactor was operated in four different conditions regarding the agitation (from 200 to 800 rpm) and the best condition was repeated to confirm the results. Results show that slowing down the rotation to 300 rpm favors riboflavin production over biomass formation (Table S4). Cultivation at this condition resulted in 2.1 g/L of riboflavin after 48h (Figure 6), corresponding to a yield of 44.2 ± 5.9 mg/L.h. The final riboflavin titer was 2.2-fold higher compared to the same strain cultivated in Erlenmeyer scale. This result shows that the sRNA-carrying strain is suitable for riboflavin production in large scale.

**Figure 6.**
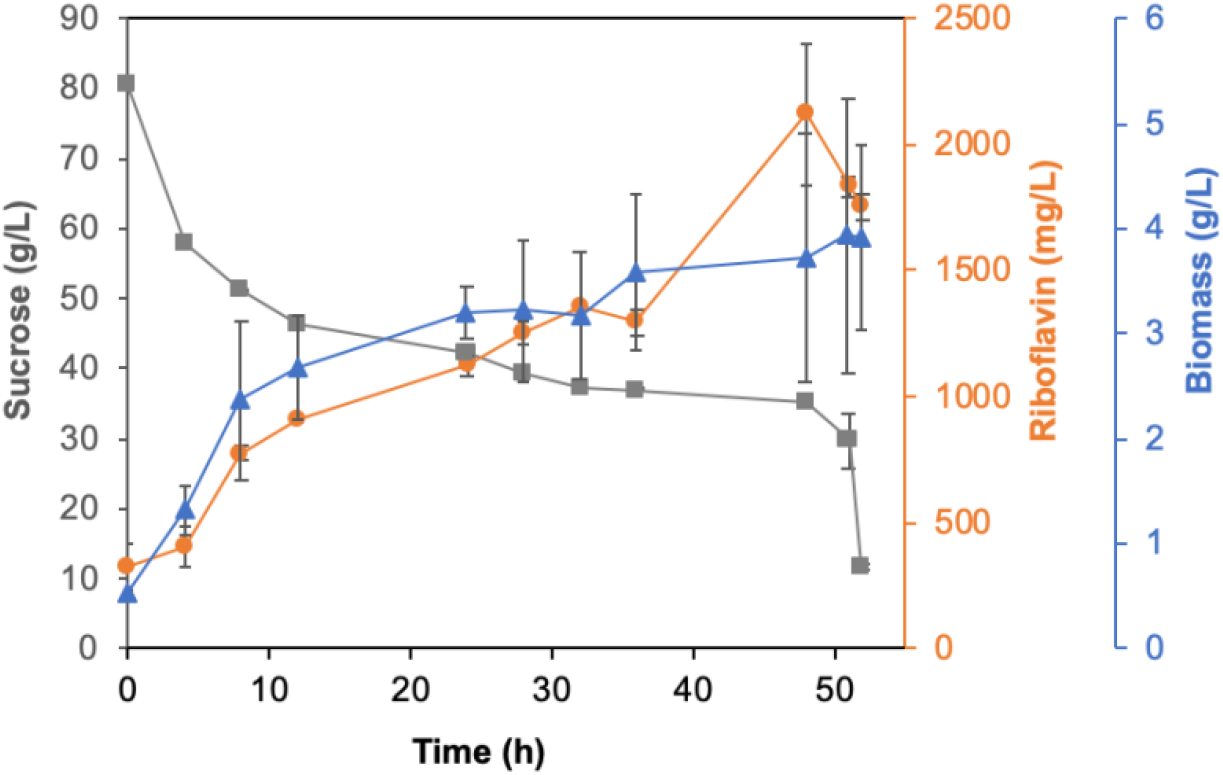
The BsRF::*ribDG*-sRNA strain is stable and highly productive in a bioreactor scale. A 5L-batch was carried out for 52h at 300 rpm, 0,5 vvm air supply, and pH controlled at 7.0±0.1. Samples were withdrawn periodically for sucrose (gray squares), riboflavin (orange circles), and biomass (blue triangles) quantification. Data are presented as mean ± SD (n = 2). Raw data available in Supplementary Data 3.

## 4. Discussion

Previously underestimated, RNA has emerged in the last decade as a versatile tool to engineer strains and regulatory circuits (Chappell et al., 2015b; Kelly et al., 2018; Leistra et al., 2019; McCarty et al., 2020). When it comes to regulating gene expression, sRNA stands up as an efficient and easy-to-engineer tool. It has been intensely used to engineer bacteria for gene knockdown by preventing translation initiation (Hoynes-O’Connor and Moon, 2016; Liu et al., 2014; Na et al., 2013; Noh et al., 2017; Yang et al., 2019). Activation of gene expression has been achieved by engineering two RNA molecules, the sRNA and a toehold switch (for translation initiation control) or a Sense target RNA (for transcription termination control) (Chappell et al., 2015a; Green et al., 2014). Our strategy differs from the latter by targeting intrinsic regulatory RNAs (riboswitches) with synthetic sRNAs to activate gene expression.

Synthetic sRNAs had been previously designed to target the riboswitches *metH, xpt*, and *pbuE*. The sRNAs targeting the *metH* and *xpt* riboswitches completely failed, and the sRNA targeting the *pbuE* riboswitch only activated gene expression by 3.1-fold (Chappell et al., 2015a). Further improvement of the *pbuE* sRNA elevated the activation to 13.4-fold (Meyer et al., 2016). We have successfully designed sRNAs targeting both the *xpt* and the *pbuE* riboswitches. Our *pbuE*-sRNA displays a 103-fold gene activation, which is much higher than achieved by others. Differently from others, our sRNA works as single genome copy. Using genome integrated sRNA genes, instead of plasmid-borne, favors strain stability for engineering metabolite producers suitable for large scale cultivation.

We used the riboswitch-targeting sRNAs to engineer a riboflavin-producing strain. The resulting strains were able to reach 50% increase in the riboflavin titer using one sRNA and 65% increase using two sRNAs. Targeting riboswitches with sRNAs seems more efficient than permanent edition. Recently, the *ribDG* riboswitch and the *purR* gene have been edited through CRISPR-Cas9 with the same goal of engineering a riboflavin-producing strain. The engineered strain produced 44% more riboflavin than the parental one (Boumezbeur et al., 2020). Interestingly, the study reports a similar improvement in culture density as observed here. Achieving similar growth improvements using different tools indicates that there are non-anticipated growth-limiting levels of gene expression within the purine and riboflavin pathways that have been alleviated by both approaches. We have developed a new synthetic sRNA tool that targets bacterial riboswitches and activates gene expression at the transcription level. The tool has been validated *in vitro* and *in vivo* against different targets demonstrating specificity. Furthermore, we used the riboswitch-targeting sRNAs to efficiently engineer a riboflavin-producing strain. Finally, we developed an easy-to-engineer multiplex sRNA to target up to five different genes simultaneously. Multiplex sRNA overcomes the obstacles faced by protein-based multiplex control. The tools we developed are broadly applicable to bacterial riboswitches, and are useful to modulate expression of essential genes that cannot be knocked out.

## Acknowledgements

This work was supported by the São Paulo Research Foundation (FAPESP) [grant 2014/17564-7 and 2020/08699-7]; Conselho Nacional de Desenvolvimento Científico e Tecnológico (CNPq) [grant 290110/2017-3 and INCT BioSyn]; and Coordenação de Aperfeiçoamento de Pessoal de Nível Superior - Brasil (CAPES) [Finance Code 001].

## Supplementary files

Supplementary material

## Mendeley data

https://data.mendeley.com/datasets/84yybmrtxx/1

DOI: 10.17632/84yybmrtxx.1

Supplementary data 1

Supplementary data 2

Supplementary data 3

