## Supplementary material for "Targeting riboswitches with synthetic small RNAs for metabolic engineering"

**Table S1.** Plasmids used or constructed in this study.

| Plasmid | Description | Reference |
|---|---|---|
| pBS1C | Integrates at the *B. subtilis amyE* locus, chloramphenicol resistance (BBa_K823023) | (Radeck et al., 2013) |
| pBS3Clux | Integrates at the *B. subtilis sacA* locus, chloramphenicol resistance (BBa_K823025) | (Radeck et al., 2013) |
| pBS2E | Integrates at the *B. subtilis lacA* locus, erythromycin resistance (BBa_K823027) | (Radeck et al., 2013) |
| pBS3C-$P_{srfA}$-*lux* | *srfA* promoter of *B. subtilis* controlling expression of *luxABCDE* | (Lins et al., 2021) |
| pBS3C-$P_{srfA}$-*purE*-*lux* | *luxABCDE* gene cluster under control of the *srfA* promoter and the *purE* riboswitch of *B. subtilis* | |
| pBS3C-$P_{srfA}$-*pbuG*-*lux* | *luxABCDE* gene cluster under control of the *srfA* promoter and the *pbuG* riboswitch of *B. subtilis* | |
| pBS3C-$P_{srfA}$-*pbuE*-*lux* | *luxABCDE* gene cluster under control of the *srfA* promoter and the *pbuE* riboswitch of *B. subtilis* | |
| pBS3C-$P_{srfA}$-*nupG*-*lux* | *luxABCDE* gene cluster under control of the *srfA* promoter and the *nupG* riboswitch of *B. subtilis* | |
| pBS3C-$P_{srfA}$-*xpt*-*lux* | *luxABCDE* gene cluster under control of the *srfA* promoter and the *xpt* riboswitch of *B. subtilis* | |
| pBS1C_ $P_{grac}$-sRNA-*purE*-broccoli | pBS1C contains *grac* promoter controlling transcription of sRNA-*purE*-broccoli | This study |
| pBS2E_$P_{grac}$-*purE*-sRNA | pBS2E contains *grac* promoter controlling transcription of sRNA designed to the *purE* riboswitch | |
| pBS2E_$P_{grac}$-RiboJ-*purE*-sRNA | pBS2E contains *grac* promoter controlling transcription of sRNA designed to *purE* riboswitch merged to RiboJ | |
| pBS2E_$P_{grac}$-*xpt*-sRNA | pBS2E contains *grac* promoter controlling transcription of sRNA designed to *xpt* riboswitch | |
| pBS2E_$P_{grac}$-*pbuE*-sRNA | pBS2E contains *grac* promoter controlling transcription of sRNA designed to *pbuE* riboswitch | |
| pBS2E_$P_{grac}$-*pbuG*-sRNA | pBS2E contains *grac* promoter controlling transcription of sRNA designed to *pbuG* riboswitch | |
| pBS2E_$P_{grac}$-*nupG*-sRNA | pBS2E contains *grac* promoter controlling transcription of sRNA designed to *nupG* riboswitch | |
| pBS2E_$P_{grac}$-*ribDG*-sRNA | pBS2E contains *grac* promoter controlling transcription of sRNA designed to *ribDG* riboswitch | |
| pBS2E_$P_{grac}$-*purR*-sRNA | pBS2E contains *grac* promoter controlling transcription of sRNA designed to *purR* gene | |
| pBS2E_$P_{grac}$-*purR*-sRNA-$P_{grac}$-*ribDG*-sRNA | pBS2E with two transcription units. 1) *grac* promoter controlling transcription of sRNA designed to *purR* gene; 2) *grac* promoter controlling transcription of sRNA designed to *ribDG* riboswitch | |
| pBS2E_$P_{grac}$-sRNA_Operon | pBS2E with six riboswitches interspaced by RiboJs, all controlled by *grac* promoter | |

**Table S2.** *B. subtilis* strains used or constructed in this study.

| Strain | Description | Reference |
|---|---|---|
| *E. coli* Top 10 | Cloning strain | Invitrogen |
| *B. subtilis* 168 | Wild-type, *trpC2* | Lab collection |
| *B. subtilis* riboflavin super-producing strain (BsRF) | P*veg* driven riboflavin gene cluster, mutations on *ribC*, *tkt* and FMN riboswitch | |
| *B. subtilis::luxABCDE* | Δ*sacA*: *luxABCDE* | (Lins et al., 2021) |
| *B. subtilis::*$P_{srfA}$-*luxABCDE* | Δ*sacA*: *srfA promoter controlling luxABCDE* | |
| *B. subtilis::*$P_{srfA}$-*purE-lux* | Δ*sacA: srfA* promoter controlling *purE* riboswitch and *luxABCDE* | |
| *B. subtilis::*$P_{srfA}$-*pbuE-lux* | Δ*sacA: srfA* promoter controlling *pbuE* riboswitch and *luxABCDE* | |
| *B. subtilis::*$P_{srfA}$-*nupG-lux* | $P_{srfA}$-*nupG-luxABCDE* integrated into the *sacA locus* | |
| *B. subtilis::*$P_{srfA}$-*pbuG-lux* | $P_{srfA}$-*pbuG-luxABCDE* integrated into the *sacA locus* | |
| *B. subtilis::*$P_{srfA}$-*xpt-lux* | $P_{srfA}$-*xpt-luxABCDE* integrated into the *sacA locus* | |
| *B. subtilis::*$P_{grac}$-sRNA-*purE*-broccoli | Δ*amyE*: *grac* promoter controlling sRNA-*purE*-broccoli transcription | This study |
| *B. subtilis::2E* | Δ*lacA*: empty vector | |
| *B. subtilis::*$P_{grac}$-*purE*-sRNA | Δ*lacA*: *grac* promoter controlling transcription of *purE*-sRNA | |
| *B. subtilis::*$P_{srfA}$-*purE-lux::*$P_{grac}$-*purE*-sRNA | Δ*sacA*: *srfA* promoter controlling controlling *purE* riboswitch and *luxABCDE*<br>Δ*lacA*: *grac* promoter controlling transcription of *purE*-sRNA | |
| *B. subtilis::*$P_{grac}$-RiboJ-*purE*-sRNA | Δ*lacA*: *grac* promoter controlling transcription of RiboJ-*purE*-sRNA | |
| *B. subtilis::*$P_{srfA}$-*purE-lux::*$P_{grac}$-RiboJ-*purE*-sRNA | Δ*sacA*: *srfA* promoter controlling controlling *purE* riboswitch and *luxABCDE*<br>Δ*lacA*: *grac* promoter controlling transcription of RiboJ-*purE*-sRNA | |
| *B. subtilis::*$P_{rsfA}$-*purE-lux::2E* | Δ*sacA*: *srfA* promoter controlling controlling *purE* riboswitch and *luxABCDE*<br>Δ*lacA*: *empty vector* | |
| *B. subtilis::*$P_{srfA}$-*purE-lux::*$P_{grac}$-*purR-sRNA* | Δ*sacA*: *srfA* promoter controlling controlling *purE* riboswitch and *luxABCDE*<br>Δ*lacA*: *grac* promoter controlling transcription of *purR*-sRNA | |
| *B. subtilis::*$P_{srfA}$-*purE-lux::*$P_{grac}$-*xpt*-sRNA | Δ*sacA*: *srfA* promoter controlling controlling *purE* riboswitch and *luxABCDE*<br>Δ*lacA*: *grac* promoter controlling transcription of *xpt*-sRNA | |
| *B. subtilis::*$P_{srfA}$-*pbuE-lux::*$P_{grac}$-*pbuE*-sRNA | Δ*sacA*: *srfA* promoter controlling controlling *pbuE* riboswitch and *luxABCDE* | |

| | |
|---|---|
| | Δ*lacA*: *grac* promoter controlling transcription of *pbuE*-sRNA |
| *B. subtilis*::$P_{grac}$-*pbuE*-sRNA | Δ*lacA*: *grac* promoter controlling transcription of *pbuE*-sRNA |
| *B. subtilis*::$P_{srfA}$-*pbuE*-*lux::2E* | Δ*sacA*: *srfA* promoter controlling controlling *pbuE* riboswitch and *luxABCDE*<br>Δ*lacA*: empty vector |
| *B. subtilis*::$P_{grac}$-*ribDG*-sRNA | Δ*lacA*: *grac* promoter controlling transcription of *ribDG*-sRNA |
| *B. subtilis*::$P_{grac}$-*purR*-sRNA | Δ*lacA*: *grac* promoter controlling transcription of *purR*-sRNA |
| *B. subtilis*::$P_{grac}$-*xpt*-sRNA | Δ*lacA*: *grac* promoter controlling transcription of *xpt*-sRNA |
| *B. subtilis*::$P_{srfA}$-*xpt*-*lux::*2E | Δ*sacA*: *srfA* promoter controlling controlling *xpt* riboswitch and *luxABCDE*<br>Δ*lacA*: empty vector |
| *B. subtilis*::$P_{srfA}$-*xpt*-*lux::*$P_{grac}$-*xpt*-sRNA | Δ*sacA*: *srfA* promoter controlling controlling *xpt* riboswitch and *luxABCDE*<br>Δ*lacA*: *grac* promoter controlling transcription of *xpt*-sRNA |
| *B. subtilis*::$P_{srfA}$-*xpt*-*lux::*$P_{grac}$-*purE*-sRNA | Δ*sacA*: *srfA* promoter controlling controlling *xpt* riboswitch and *luxABCDE*<br>Δ*lacA*: *grac* promoter controlling transcription of *purE*-sRNA |
| *B. subtilis*::$P_{srfA}$-*xpt*-*lux::*$P_{grac}$-RiboJ-*purE*-sRNA | Δ*sacA*: *srfA* promoter controlling controlling *xpt* riboswitch and *luxABCDE*<br>Δ*lacA*: *grac* promoter controlling transcription of RiboJ-*purE*-sRNA |
| *B. subtilis*::*P*$_{srfA}$-*xpt*-*lux::P*$_{grac}$- *purR*-sRNA | Δ*sacA*: *srfA* promoter controlling controlling *xpt* riboswitch and *luxABCDE*<br>Δ*lacA*: *grac* promoter controlling transcription of *purR*-sRNA |
| BsRF*::purR-sRNA-ribDG*-sRNA | Δ*lacA*: *grac* promoter controlling transcription of *purR*-sRNA and a second *grac* promoter controlling transcription of *ribDG*-sRNA |
| BsRF*::* sRNA_Operon | Δ*lacA*: operon of six riboswitches interspaced by synthetic RiboJs |

**Table S3. sRNA design parameters**

| target | ΔG Kcal/mol[a] | | | | | equilibrium |
|---|---|---|---|---|---|---|
| | aptamer | sRNA | seed | hybridization | complex | |
| *purE* riboswitch | -18.8 | -1.7 | -7.2 | -21.8 | -58.8 | 100% |
| *pbuE* riboswitch | -10.2 | -2.6 | -7.2 | -21.2 | -57.2 | 100% |
| *xpt* riboswitch | -17.6 | -6.1 | -8.4 | -22.4 | -59.4 | 98% |
| *pbuG* riboswitch | -18.0 | -2.6 | -7.2 | -20.3 | -62.1 | 100% |
| *nupG* riboswitch | -21.5 | -7.6 | -4.8 | -21.5 | -58.7 | 100% |
| *purR* gene | na | -1.1 | na | -52.3 | -109.8 | 100% |

[a] at 37ºC

[b] complex representation at equilibrium

na = not applicable

The seed energy is given by the free energy release due to the interaction between the seed regions (unpaired nucleotides that initiate the hybridization). The hybridization energy is the free energy release due to the mRNA-sRNA base-pairing. The energy of the complex is the free energy release due to the mRNA-sRNA complex formation.

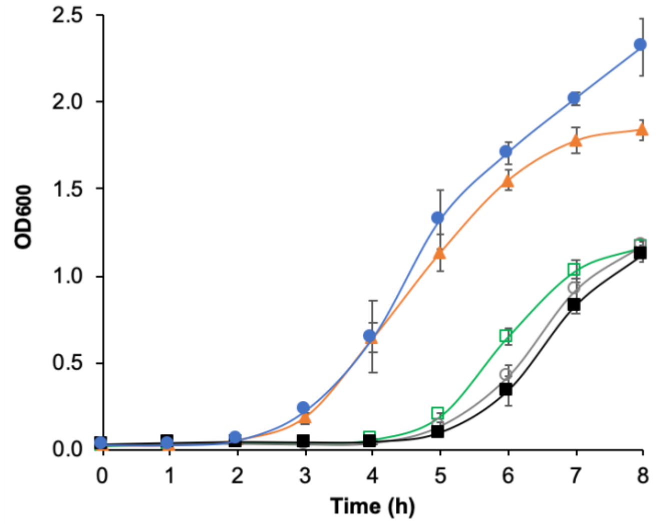


**Figure S1.** Growth curve of *B. subtilis* strains carrying the lux operon controlled by a riboswitch only or the later and a sRNA. *B. subtilis*::*pbuE* riboswitch-lux (green open square), *B. subtilis*::*pbuE* riboswitch-lux::*pbuE*-sRNA (orange filled triangle), *B. subtilis*::*purE* riboswitch-lux (gray open circle), *B. subtilis*::*purE* riboswitch-lux::RiboJ-*purE*-sRNA (blue filled circle), *B. subtilis*::*purE* riboswitch-lux::*purE*-sRNA (black filled square).

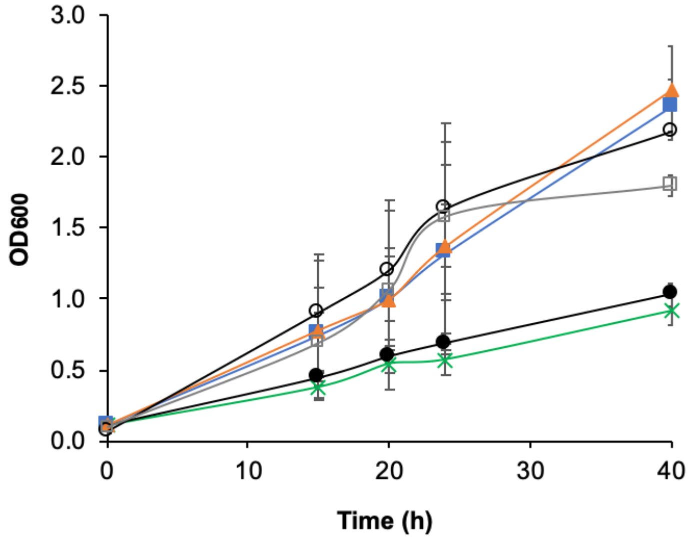


**Figure S2.** Growth curve of *B. subtilis* riboflavin producing strains (BsRF). BsRF parental strain (green x), BsRF::*purR*-sRNA (orange closed triangle), BsRF::*ribDG*-sRNA (blue closed square), BsRF::*xpt*-sRNA (black closed circle), BsRF::*ribDG*-sRNA-*purR*-sRNA (black open circle), BsRF::sRNA-Operon (gray open square).

**Table S4**. Riboflavin production by the BsRF::*ribDG*-sRNA strain in bioreactor 5L-batch under different agitation rates.

| Agitation (rpm) | Biomass ($g.L^{-1}$) | Riboflavin ($g.L^{-1}$) | $\mu_{max}$ ($h^{-1}$) | $Y_{p/x}$ ($g.g^{-1}$) | $Y_{x/s}$ ($g.g^{-1}$) | Yield ($mg.L^{-1}.h^{-1}$) |
|---|---|---|---|---|---|---|
| 800 | 7.2 | 0.69 | 0.174 | 0.093 | 0.105 | 49.0 |
| 400 | 5.8 | 1.28 | 0.171 | 0.240 | 0.102 | 60.9 |
| 300 | 3.9 | 2.12 | 0.089 | 0.655 | 0.047 | 44.2 |
| 200 | 2.0 | 0.72 | 0.148 | 0.444 | 0.011 | 24.0 |

**DNA sequence of P$_{grac}$_sRNA-*purE*-broccoli**

GGTACCAGCTATTGTAACATAATCGGTACGGGGGTGAAAAAGCTAACGGAAAAGGGAGCGGAAAAGA
ATGATGTAAGCGTGAAAAATTTTTTATCTTATCACTTGAAATTGGAAGGGAGATTCTTTATTATAAGAAT
TGTGGAATTGTGAGCGGATAACAATTCCCAATTGATTCAACAATTATGTGAGGTCGTGTTTCTTTGCCAT
GTGTATGTGGGAGACGGTTCCAGATATTCGTATCTGTCGAGTAGAGTGTGGGCTCCCACATACTCTGAT
GATCCTTCGGGATCATTCATGGCAACAAAGCCCGCCGAAAGGCGGGCTTTTCTGT

P$_{grac}$

*purE*-sRNA

RNAbroccoli

T500 terminator

**DNA sequence of the sRNA Operon**

GGTACCAGCTATTGTAACATAATCGGTACGGGGGTGAAAAAGCTAACGGAAAAGGGAGCGGAAAAGA
ATGATGTAAGCGTGAAAAATTTTTTATCTTATCACTTGAAATTGGAAGGGAGATTCTTTATTATAAGAAT
TGTGGAATTGTGAGCGGATAACAATTCCCAATTAGCTGTCACCGGATGTGCTTTCCGGTCTGATGAGTC
CGTGAGGACGAAACAGCCTCTACAAATAATTTTGTTTAAGATTCAACAATTATGTGAGGTCGTGTTTCTA
GTCGTCAAGTGCTGTGCTTGCACTTCTGATGAGGCAGTGATGCCGAAACGACCTCTACAAATAATTTTG
TTTAACGACGAAACTTCATGAACTCACCCCCAGTCGTCAAGTGCTGTGCTTGCACTTCTGATGAGGCAG
TGATGCCGAAACGACCTCTACAAATAATTTTGTTTAAGAAATTAATACGACTCACTATAGGGCCTATCCT
GGCTCCGAAGGGTGCATATTCAAGCGCTCAACGGGTGTGCTTCCCGTTCTGATGAGTCCGTGAGGACG
AAAGCGCCTCTACAAATAATTTTGTTTAACCATATCTACGATGAGAGTTGTGTTCCAAGAGGAGTCAATT
AATGTGCTTTTAATTCTGATGAGACGGTGACGTCGAAACTCCCTCTACAAATAATTTTGTTTAACCATAG
GTTATACAAGTGATGAGAGACGCCAGCTGTCACCGGATGTGCTTTCCGGTCTGATGAGTCCGTGAGGA
CGAAACAGCCTCTACAAATAATTTTGTTTAACAAAGCCCGCCGAAAGGCGGGCTTTTCTGT

P$_{grac}$ promoter

*purE*-sRNA

*purR*-sRNA

*ribDG*-sRNA

*xpt*-sRNA

*pbuE*-sRNA

RiboJ 60

RiboJ

RiboJ10

RiboJ64

T500 terminator
